# Ongoing worldwide homogenization of human pathogens

**DOI:** 10.1101/009977

**Authors:** T Poisot, C Nunn, S Morand

## Abstract

**Background:** Infectious diseases are a major burden on human population, especially in low- and middle-income countries. The increase in the rate of emergence of infectious outbreaks necessitates a better understanding of the worldwide distribution of diseases through space and time.

**Methods:** We analyze 100 years of records of diseases occurrence worldwide. We use a graph-theoretical approach to characterize the worldwide structure of human infectious diseases, and its dynamics over the Twentieth Century.

**Findings:** Since the 1960s, there is a clear homogenizing of human pathogens worldwide, with most diseases expanding their geographical area. The occurrence network of human pathogens becomes markedly more connected, and less modular.

**Interpretation:** Human infectious diseases are steadily expanding their ranges since the 1960s, and disease occurrence has become more homogenized at a global scale. Our findings emphasize the need for international collaboration in designing policies for the prevention of outbreaks.

**Funding:** T.P. is funded by a FRQNT-PBEE post-doctoral fellowship, and through a Marsden grant from the Royal Academy of Sciences of New-Zealand. Funders had no input in any part of the study.

## Introduction

Transmissible infectious diseases impose a major burden on human populations worldwide, being accountable for thirteen millions deaths a year. Fifty percent of these deaths occur in developing countries, representing up to 80% of all age-normalised deaths in some countries. The composition of pathogen communities has far-reaching impacts on political stability, economics, and human behaviour [1]. The frequency and severity of outbreaks may also accelerate as human pressures on ecosystems become stronger [2], and as urban areas with high population density and contact rates became larger. This may be especially true in developing countries where the majority of the population lacks access to adequate health care. Even developed countries are put at a significantly heightened risk of outbreaks when the economy is destabilized [3–5]. This can lead in the re-emergence of diseases though to be locally eradicated [6]. Since the 1960s, the number of countries reporting outbreaks follows an exponential increase, as does the number of pathogens involved in outbreaks [7]. Most efforts to address the rising challenge of increased outbreaks (both in the number of events and number of pathogens involved) have been country-centered [8]; future efforts should adopt a global perspective for at least two reasons. First, countries are not independent units, and those sharing a high proportion of their pathogens should be managed in similar ways. Second, the rising number of epidemic events means that the biogeographic distribution of pathogens is likely changing as pathogens increase their geographic range, and requires a global rather than local approach to be efficiently managed.

Previous studies focused on classical epidemiological measures, such as richness or prevalence, and correlated these measures to local climatic and diversity variables. This correlative approach yielded important results, notably the existence of a correlation between local diversity of reservoir species and disease prevalence and a latitudinal gradient in pathogen richness [8]. However, this approach mostly considers that global trends emerge from local environmental variables or socio-economical conditions.

Another perspective on the distribution of human pathogens is to explicitly consider the interactions between pathogens and countries at the global scale, and to correlate the position of countries in this complex system in relation to their epidemiological properties. Global approaches have proven fruitful to understand the worldwide dynamics of a single pathogen [9], but are yet to be applied on the ensemble of all human pathogens. Network-based approaches resulted in major breakthroughs in epidemiology, such as a better understanding of pathogen spread within complex communities [10,11] or formal rules for disease spread management [12]. Network approaches also hold promise to address this global challenge.

Here we use a network-based approach to analyse changes in the biogeographic structure of human pathogens since the 1910s. Site-species (here country-pathogen) occurrence matrices are a powerful representation of communities with a biogeographic structure [13]. Using a sliding-window analysis, we discovered evidence for a worldwide homogenization of pathogens with steadily increasing biogeographic ranges.

## Methods

### Data selection and preparation

We used data from the GIDEON (*Global Infectious Diseases and Epidemiology Network*, www.gideononline.com) database to reconstruct a bipartite network of human pathogens occurrences in countries. We subset this database by bins of ten consecutive years (so that a bin for year *Y* covers all years from *Y* to *Y +*9), starting from the earliest date, and going to the farthest, with steps of one year. Each data point is a summary of ten years of human pathogens occurrences worldwide. Binning enables us to compensate for incomplete, missing, or delayed reporting of outbreaks. Although modern reporting systems in developing countries have excellent response times [14], it is likely that the records for the first half of the twentieth century, or for emerging countries, are more fragmented.

For each ten-years bin, we reconstruct the bipartite network *𝒫*_*t*_(C;P) describing the occurrence of pathogens P in countries C in the following way: each country and each pathogen which are mentioned at least once are nodes of the network; an edge exists between a country and a pathogen if there is at least one recorded occurrence of this pathogen in this country over the ten-years period.

### Network metrics

Within each bin, we apply the following network structure metrics. First, we measure the network density,

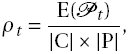

where |C| and |P| are, respectively, the number of countries and pathogens in the bin, and E(*P*_*t*_) is the total number of outbreaks recorded (outbreaks of the same pathogen in the same countries count only once). Values of *ρ*_*t*_ range from approx. 0 to 1; values close to 0 indicate that each country has a single, unique pathogen, and values close to 1 indicate that all pathogens occured in all countries during the bin.

Second, we measured the *Nestedness based on Overlap and Decreasing Fill* algorithm to estimate the biogeographic structure of the network [15]. NODF measures the extent to which (1) countries with a large number of occurences have pathogens from countries with fewer occurences, and (2) pathogens found in countries with few occurences are also found in countries with more occurrences. NODF ranges from 0 to 100, with values close to 0 indicating that each country has its own set of pathogens, and values close to 100 indicating that the pathogen-poor countries have a subset of the pathogens from the pathogen-rich countries.

Finally, we measure the bipartite network modularity *𝒬* [16], optimized through the LP-BRIM method [17]. Modularity reflects the fact that some groups of vertices interact more together than with vertices from other groups (here meaning that there are groups of countries, each sharing a distinct set of pathogens). *2* is measured as

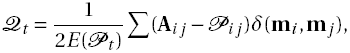

where **A** is the incidence matrix of *𝒫*_*t*_, **P** is a matrix giving the probability that pathogen *j* is found in country *i* during the bin (which is proportional to the number of countries that *i* is found in, and the number of pathogens in *j*). **m** is a vector giving the identity of the module to which each country and pathogen belong, and *δ* is Krönecker’s function, which takes a value of 1 if its two parameters are equal, and 0 otherwise. The LP-BRIM algorithm transmits random labels over edges in the network until the value of *Q* reaches a minimum, after what it reassigns the nodes increasing modularity the most to other modules, until an optimum is found. This method is known to give fast and optimal results in networks or moderate to large size.

The LP-BRIM method returns the optimized value of *𝒫*, with values close to 1 indicating that edges are established only between vertices frome the same module (*i.e.* there exists groups of countries that share no pathogens between them), and values close to 0 denoting a lack of modularity. The method also returns the community partition **m**, *i.e.* the module to which each country/pathogen belongs, and the total number of such modules.

## Results

### Number of pathogens and their occurrence increased in the last 100 years

In *Figure 1*, we represent the number of pathogens with a least one recorded worldwide occurrence in the ten-year period, and the total number of pathogen-country pairs observed. As reported by Reference 7, the *raw* number of outbreaks is higher than that, and increases exponentially since the 1960s.

**Figure 1:**
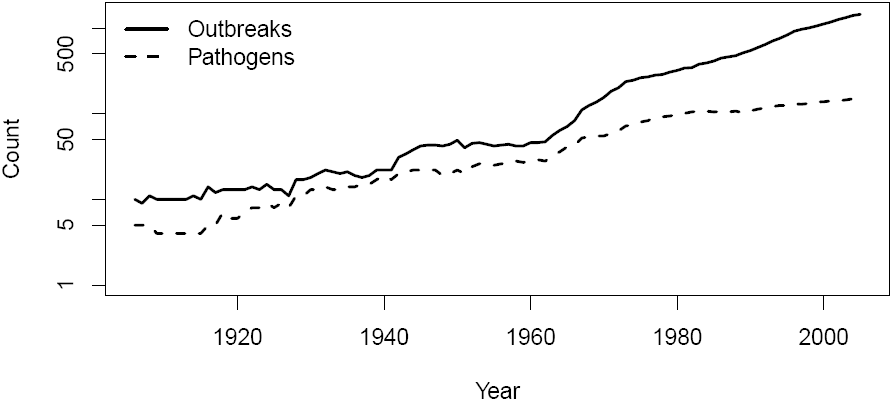
Raw number of outbreaks and pathogens over time.

### The density of the country-pathogen network is increasing since the 1980s

The network density went from 0.0374 in 1967 to 0.0651 in 2005 (*Figure 2*). This means that on average, outbreaks of a given pathogen occurred in twice as many countries in the last 10 years than they did 40 years before. This apparently low increase must be viewed in the fact that network density is a multiplicative process, and that for an increase in density when the network size (number of pathogens times number of countries) increases indicates that pathogens occurrences are more frequent.

**Figure 2:**
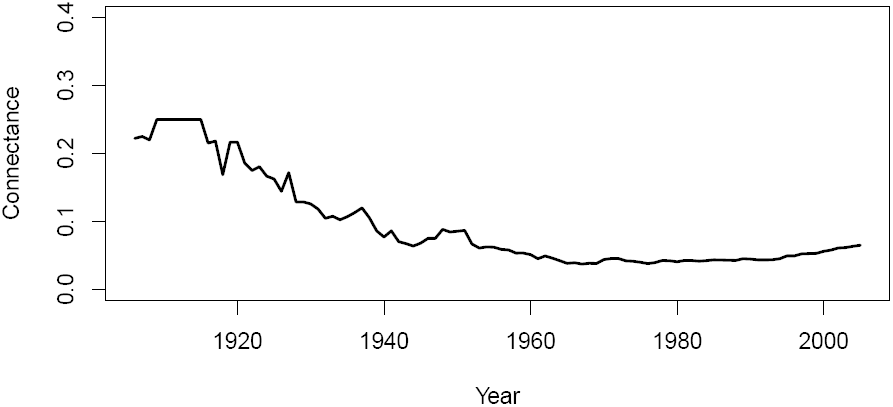
Temporal dynamics of the pathogen–country network.

### The number of modules is decreasing since the 1970s

The number of modules (groups of countries and their pathogens) reached a maximal value of 21 in 1965 (*Figure 3*). This reflects a situation in which human pathogens were highly structured from a biogeographic point of view. Yet since this period, the number of modules is rapidly decreasing, at the rate of approximately one module lost every three years. Over the period from 1960 to 2000, the correlation between the number of modules and time is markedly negative (-0.83; df=38; t=-6.58).

**Figure 3:**
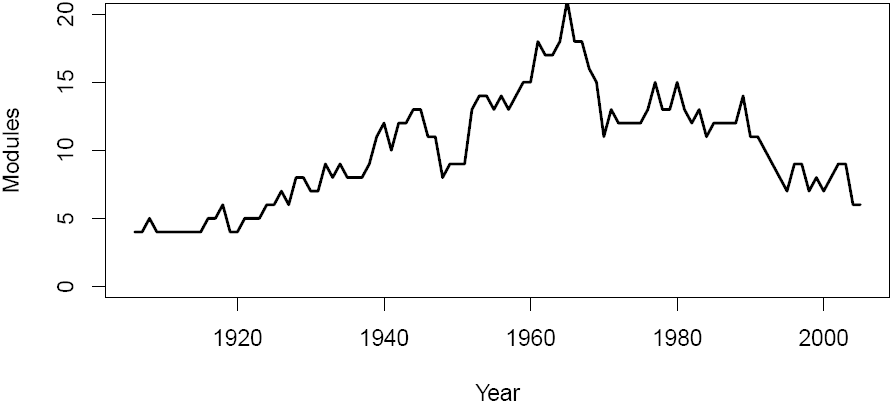
Number of modules in the pathogen–country network. Seeing as the the number of pathogens increases, the increase in connectance since the 1970s is a sign of a fast increase in the number of countries where these pathogens are found.

### The modularity is decreasing since the 1960s

(*Figure 4*). Modularity was stable in the dataset prior to the 1960s, with values close to 0.7, indicative of a high modularity (*i.e.* high biogeographic structure). The peak modularity (*2* = 0.914) was reached in 1961. Since this period, the modularity has been decreasing in a linear way (from 1960 to 2000, -0.97; df=38; t=-20.01), reaching a value of 0.2981 in 2005. Previous analyses [13,18] indicate that values of modularity around 0.3 cease to be statistically significant.

**Figure 4:**
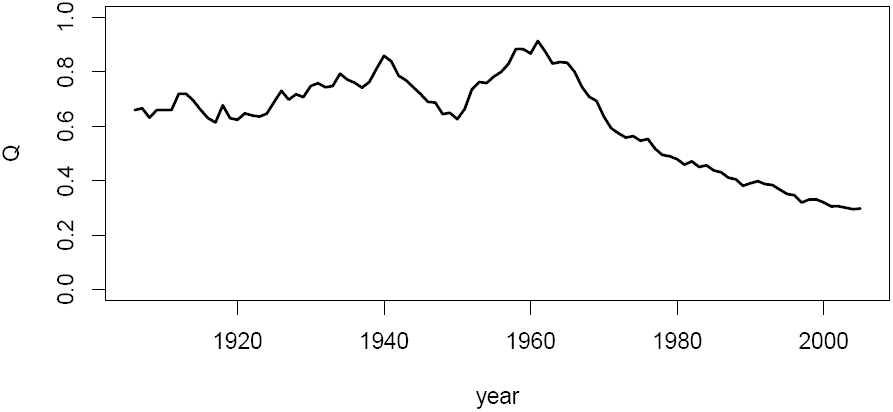
Temporal dynamics of the pathogen–country modularity. The decrease in modularity starts within 5 years from 1960.

## Discussion

We demonstrated that human pathogens have consistently increased their worldwide distribution. This increase has not happened only through migration into neighboring countries, but is rather consistent with a blurring of the area of distribution of pathogens, than have increased opportunities to cross several national boundaries at once. Although the number of pathogens has kept approximately constant in recent decades, individual pathogens increased their ranges. This results in a striking decrease of the modularity of the country-pathogen network; this is suggestive of a loss of biogeographic structure of pathogen distributions, or in other words a global homogenization of human pathogens. This homogenization occurs at the worldwide scale and has shown no signs of slowing down for the last 50 years, indicating that internationally coordinated and efficient health policies are desperately needed. The current Ebola virus outbreak exemplifies this necessity: the virus has been brought into the continental US [19] onboard a commercial flight from Liberia; the virus managed to cross several national boundaries across three continents before the necessary international effort was invested to control it. Managing the spread of pathogens at the global scale cannot be made without accounting for the dispersal of infected individuals.

This rapid decrease in modularity is opposite to network Dynamics in the first one-half of the 20th Century, which showed an increase of both modularity and the number of different modules up to the 1960s. One of the most plausible causes for the pre-1960s dynamics is methodological. Better disease surveillance and detection should result in increased and accelerated reporting of individual outbreaks. These modules correspond to newly discovered or reported pathogens in countries that have not been extensively studied before.

However, the constant changes in modularity and other aspects of network structure since the 1960s happened despite the fact that the number of countries and pathogens reported were more or less constant. This means that these changes are driven by changes in the biogeographical distribution of pathogens. Multiple factors are likely involved with these changing biogeographical distributions. Air traffic [20], marine transport [21], and all other forms of human movements across national boundaries at the global scale [22] reduce constraints on pathogen movement, and are the most likely contributors to the global homogenization that we quantified in this study.

To conclude, we used network approaches to reveal a long-term, worldwide change in the biogeographic structure of human pathogens. These results indicate that pathogens are spreading across national boundaries at a steady rate; they make intuitive sense, and are of high relevance for policy making, yet had not been previously been demonstrated. This bears important consequences for the management of pathogen spread. First, local policies are unlikely to have high success, as they cannot account for the global movement of pathogens. Second, management actions should pay attention to countries that act as sources of new pathogens, especially those countries with high movement of people and goods internationally. Additionally, well-connected *hub* countries could be preventively targeted, as they are in a position to limit the international flow of pathogens. Finally, in line with the previous idea, it is important to identify the specific pathways of disease movement across countries, and to develop effective ways to monitor these movements.

